# Abnormally high levels of serum α-klotho result in a poor outcome for clinical pregnancy-A prospective cohort study

**DOI:** 10.1101/113936

**Authors:** Miyako Funabiki, Sagiri Taguchi, Yoshitaka Nakamura

## Abstract

**Background:** The klotho protein has been extensively studied. However, there are no studies examining the association between serum alpha klotho levels and the clinical outcome of post-clinical pregnancy.

**Methods:** We conducted a prospective cohort study in 42 patients (median age 37.4 years) to evaluate the association between serum alpha klotho levels during the follicular phase of preimplantation and the clinical outcome data of post-clinical pregnancy. The patients provided informed consent at our clinic. The serum alpha klotho levels were evaluated using a human soluble alpha klotho assay kit. The fetal chromosomal abnormalities were investigated at our clinic. We also assessed the clinical outcomes of post-clinical pregnancies.

**Results:** The serum alpha klotho level during the follicular phase of preimplantation for non-pregnant women was 544.31 pg/ml (mean). The clinical pregnancy rate was 38.1%. There were chromosomal abnormalities observed in four unborn children (9.5%; Down syndrome, etc). The serum alpha klotho levels during the follicular phase of preimplantation in the chromosomal abnormalities group were higher than in the group without chromosomal abnormalities (P=0.029, abnormalities group 659.26 pg/ml [mean] versus control 530.23 pg/ml [mean]). A multiple logistic regression analysis showed the chromosomal abnormalities rates in unborn children were positively influenced by serum alpha klotho levels during the follicular phase of preimplantation (p=0.0008) and the patient’s age (p=0.008).

**Conclusion:** Previous studies have demonstrated that increased alpha klotho levels in human serum are positively correlated with health. However, abnormally high levels of serum alpha klotho during the follicular phase of preimplantation may predict a poor outcome for clinical pregnancy.

## Introduction

Klotho protein was detected in 1997 ^1^. Although klotho is intensively researched, α-klotho is known as an anti-aging molecule. Furthermore, increases in α-klotho concentrations in human serum positively promote human health ^2, 3^. Therefore, increases in α-klotho concentrations in human serum positively may improve the clinical pregnancy rates and the clinical outcome of post-clinical pregnancy. However, there are no prospective cohort studies examining the association between serum α- klotho levels and the clinical outcome of post-clinical pregnancy.

## Methods

### Study design

We conducted a prospective cohort study in 42 patients (median age 37.4 years) to evaluate the association between serum α-klotho levels during the follicular phase of preimplantation and the clinical outcome data of post-clinical pregnancy at our clinic from May 2015 to September 2016. Furthermore, our prospective cohort study was also conducted according to STROBE guideline for cohort study reporting.

### Institutional Review Board (IRB) approval

Our study was approved by the IRB of Oak Clinic, Japan. The patients provided informed consent at our clinic.

### Human soluble α-klotho assay

Human blood samples were drawn from a forearm vein in the morning after overnight fasting. Sera were obtained by centrifugation and immediately stored at −30 °C. The serum α-klotho levels were evaluated using a human soluble α-klotho assay kit (TAKARA BIO Inc., Japan).

### The investigation for the fetal chromosomal abnormalities

The fetal chromosomal abnormalities were investigated by physicians at our clinic. We also assessed the clinical outcomes of post-clinical pregnancies.

### Statistical tests

The statistical tests were performed using Dr. SPSS II for Windows (SPSS Japan, Inc., Tokyo), and significance was defined as p < 0.05 (two-tailed). Statistical analyses were investigated by using *Welch's* t-*test*, Spearman's rank-correlaion test and multiple logistic regression analysis.

## Results

### Results 1

The serum α-klotho level during the follicular phase of preimplantation for non-pregnant women was 544.31 pg/ml (mean). The clinical pregnancy rate was 38.1%.

There were chromosomal abnormalities observed in four unborn children (9.5%; Down syndrome, *etc:* Figure 1).

**Figure. 1.**
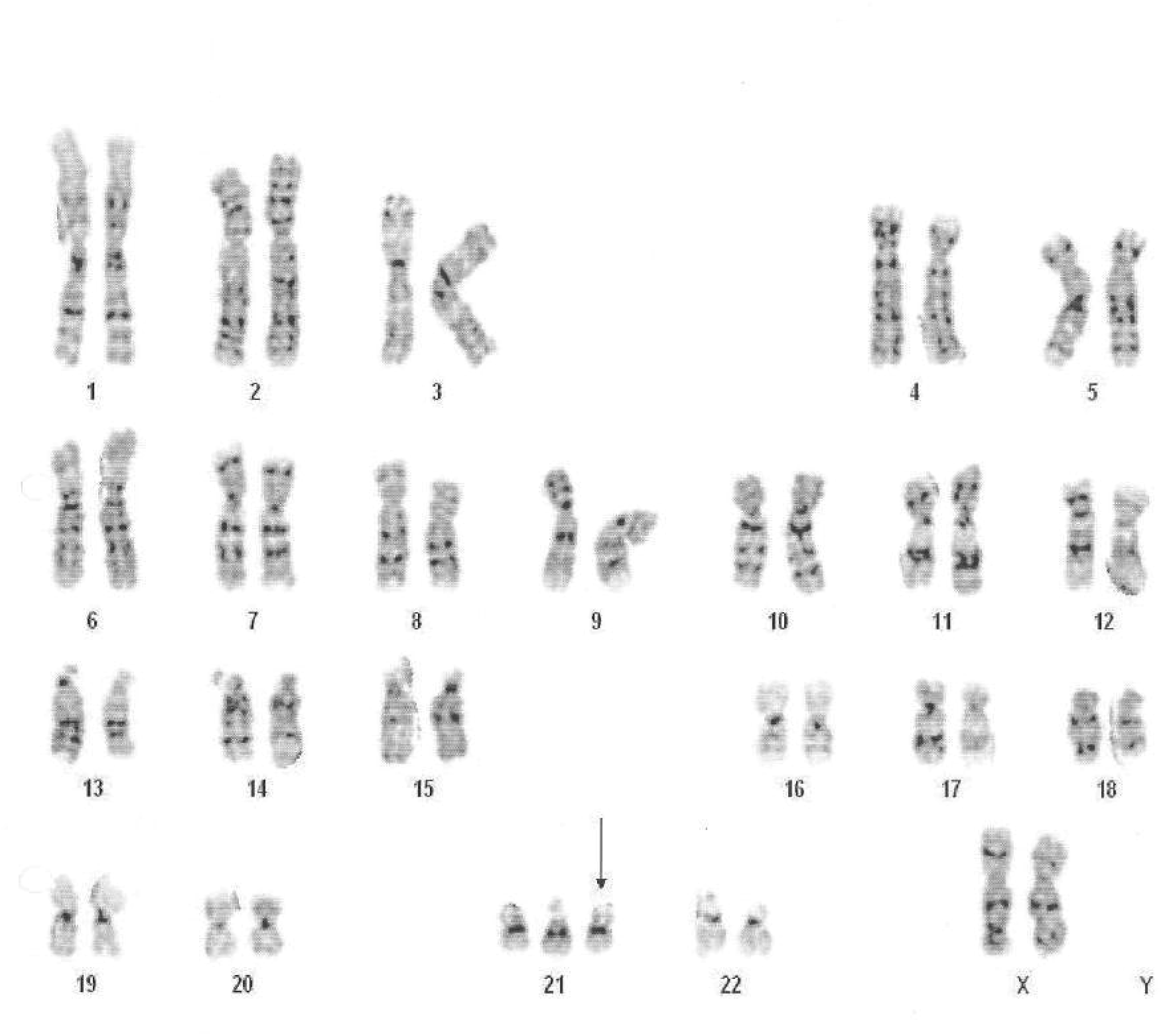
A typical image (Down syndrome case) for chromosomal abnormalities observed in unborn children

### Results 2

The serum a α-klotho levels during preimplantation (r= - 0.496, P<0.01) and the pregnancy rates (r= - 0.337, P<0.01) were decreased by aging (Table 1), while the pregnancy rates were improved by increasing of the serum α-klotho levels during preimplantation.

**Table 1:**
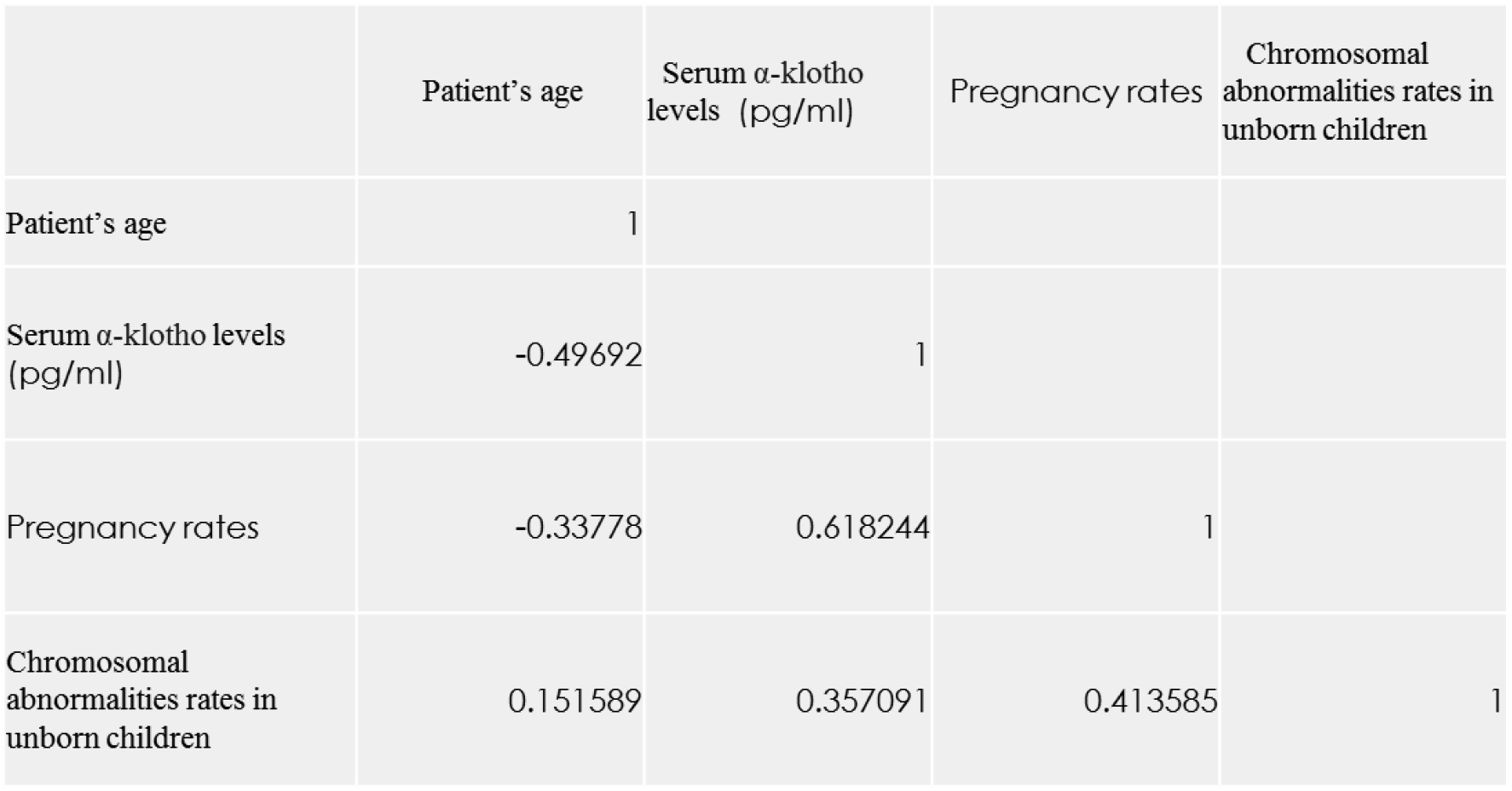
Spearman's rank-correlation coefficient

However, the chromosomal abnormalities rates in unborn children were observed by increasing of the serum α-klotho levels during preimplantation (r= 0.357, P<0.01: Table 1).

### Results 3

The serum a α-klotho levels during the follicular phase of preimplantation in the chromosomal abnormalities group were higher than in the group without chromosomal abnormalities (P=0.029; abnormalities group 659.26 pg/ml [mean] versus control 530.23 pg/ml [mean]).

### Results 4

A multiple logistic regression analysis (Table 2) showed the chromosomal abnormalities rates in unborn children were positively influenced by serum α-klotho levels during the follicular phase of preimplantation (p=0.0008) and the patient’s age (p=0.008).

**Table 2:** Multivariate logistic regression analysis 95% CI means 95 *%* confidence interval.

|  | T value | P value | 95% CI (Lower) | 95% CI (Upper) |
| --- | --- | --- | --- | --- |
| Patient's age | 2.773825 | 0.008457 | 0.008909 | 0.056891 |
| Serum α-klotho levels (pg/ml) | 3.645358 | 0.000778 | 0.000618 | 0.002157 |

## Discussion

Previous studies have demonstrated that increased a α-klotho levels in human serum are positively correlated with health ^2. 3^. However, we abnormally high levels of serum α-klotho during the follicular phase of preimplantation may predict a poor outcome for clinical pregnancy. Therefore, an abnormal increase in a α-klotho is not always beneficial for humans. Although we speculate that the abnormally high levels of serum α-klotho during the follicular phase of preimplantation could be directly related to the chromosomal abnormality of her unborn child, investigation of the detailed mechanism by the use of next-generation sequencing (NGS) technology would be necessary in the near future.

## Author contributions

M.F, S.T. and Y.N.: Concept, Provision of the study materials, collection and/or assembly of the data, analysis and interpretation of the data and final approval of the manuscript.

## Acknowledgements

We are grateful to the physicians, nurses and clinical embryologists for their assistance with the design of this study and/or the experiments performed at our clinic.

